# Nano-computed tomography reveals repeated phenotypic divergence in parasites to escape host defense

**DOI:** 10.1101/2023.01.21.525028

**Authors:** Stanislav Kolencik, Edward L. Stanley, Aswaj Punnath, Avery R. Grant, Jorge Doña, Kevin P. Johnson, Julie M. Allen

**Affiliations:** Department of Biology, University of Nevada Reno, Reno, NV, U.S.A.; Department of Natural History, Florida Museum of Natural History, University of Florida, Gainesville, FL, U.S.A.; Entomology and Nematology Department, University of Florida, Gainesville, FL, U.S.A; Illinois Natural History Survey, Prairie Research Institute, University of Illinois at Urbana Champaign, Champaign, IL, U.S.A.; Departamento de Biología Animal, Universidad de Granada, Granada, Spain

## Abstract

Understanding how selective pressures drive morphological change is a central question in evolutionary biology. Feather lice have repeatedly diversified into convergent ecomorphs, based on how they escape from host defenses in different microhabitats. Here, we used nano-computed tomography scan data of 89 specimens of feather lice, belonging to four ecomorph groups to quantify variation of functional traits, including mandibular muscle volume, limb length, and three-dimensional head shape data in these tiny insects. The results suggest that the shape of the head, the proportional volume of the chewing muscles, and the length of the leg segments in feather lice are all strongly associated with specific host-habitats. Further, species that co-occur on hosts have increased rates of morphological evolution, suggesting competition for host space is one of the drivers of morphology. This supports previous work indicating that the phenotypic diversity of feather lice is the result of repeated convergence resulting from resource partitioning, microhabitat specialization, and selection pressures imposed by host defense.

## INTRODUCTION

The evolutionary trajectory of an organism is shaped by its ecology. However, understanding the link between broad macroevolutionary patterns and ecological processes has been incredibly challenging. Organisms that have undergone similar evolutionary shifts (convergence) provide excellent models to examine hypotheses about these links. Systems with a high degree of isolation, such as islands or lakes, seem particularly efficient engines of diversification and morphological convergence [e.g., limb proportions in Anolis lizards (*1*), jaw morphology in African rift lake cichlids (*2*), and beak shape in Hawaiian honeycreepers (*3*)]. For permanent parasites, hosts are analogous to islands (*4*) because of limited dispersal between host species. In these systems, parasites often specialize either by using a particular microhabitat of the host (*5*) or in how they escape from host defenses (*6*). However, the traits that characterize these radiations are not well understood, due to the challenges of quantifying 3D morphology in microscopic organisms.

Avian feather lice are a group of permanent ectoparasites that show evidence of repeated convergence in ecomorphological features in response to host defense (*7*). Because lice spend their entire life cycle on a single host and do not fly or jump, they often have limited dispersal opportunities to other species of birds, which might promote in situ diversification (*6, 8*). Some birds are host to multiple species of lice (*9*), and competition between lice could result in resource partitioning and subsequent microhabitat specialization. A high louse infestation rate can cause significant feather damage resulting in lowered ability to thermoregulate and may result in secondary infections and symptoms such as restlessness, anorexia, weight loss, reduced egg laying, and even the death of the infected bird (*9, 10*). Therefore, birds and lice are trapped in a coevolutionary system where birds have evolved defenses to remove ectoparasites; conversely, lice have evolved counter adaptations to host defenses to remain attached to the host.

Birds have two main defenses against lice: preening with the bill and scratching with the feet (*11*). Feather lice (Phthiraptera: Ischnocera) have evolved into different ecomorphs depending on the part of the host’s body they occupy (*7, 12–14*) and how they escape from host defenses. The most common ecomorphs found across feather lice are termed wing, head, body, and generalist lice (*7, 12*). Wing lice are long and slender in shape and insert between barbs of the bird’s wing feathers, making it difficult for the bird to remove them through preening (*15*). Head lice have a plump, rounded body with a triangular head, likely supporting strong mandibular muscularization. These lice remain on the head where the bird cannot preen with its bill, and it is thought they use this muscularization to bite down and hold onto feather barbs to avoid being removed by scratching (*12, 16*). Body lice have a short, rounded body and head, and escape host defenses by burrowing into the downy parts of the feathers on the body of the bird (*17*). Some feather lice are more generalized in their body form and can be found over many regions of the host (*12*). Categorization of the microhabitat specialization of lice has been based on observational evidence (*12, 13*) and qualitative assessments of general morphological form (*14*). Previous phylogenetic analysis, based on three loci (*7*), found that each ecomorph has evolved repeatedly. These results indicate there has been repeated morphological divergence within host groups and convergent evolution across feather louse lineages. However, a detailed study characterizing and quantifying the morphological features specific to each ecomorph and analyzing their differences has not been done. Quantitative assessments of functional traits have some advantages over a purely categorical approach (*18*) as they allow a more nuanced assessment of morphological disparity.

Traditional morphological analyses of lice use slide-mounted specimens and microscopy to find morphological differences between species and to make detailed descriptions of new species (e.g., *19*). This method is advantageous as it is a relatively simple process, requiring only a slide-mounted specimen and a microscope with an ocular scale. This method can identify the presence or absence of some characteristic structures and recover two-dimensional measurements, such as length and width, but is limited when taking linear measurements in a complex three-dimensional environment (e.g., depth) and cannot recover volumetric data. This gap can be filled by using nano-computed tomography (CT). This method uses multiple X-ray images to reconstruct cross sections of an object, which can be used to produce 3-D visualization of the external and internal structures allowing for quantitative measurements and observations.

Here, we collected and analyzed nano-CT scan data from different louse species representing several independent evolutionary origins of ecomorph (head, body, wing, and generalist). We investigated the hypothesis that convergence in the overall morphology of feather louse ecomorphs results from the same specific morphological transitions. We focused on two features that are believed to be adaptive for each ecomorph in escaping host defense. First, it has been hypothesized that the main counter-defense of head lice to avoid being removed by scratching is to slide a feather barb into the cranial rostral groove and bite with the mandibles to remain attached. Therefore, large mandibular muscles might be favored and could be the reason for convergent head lice to have an expanded temple region as compared to other ecomorphs. Therefore, we predict head lice will have the largest mandibular muscularization. We quantified this by calculating the proportional volume of the chewing muscles and using a geometric morphometric approach to recover three-dimensional head shape variation across ecomorphs. Second, we predict differences in proportional lengths of the leg parts (tibia and femur), as they may vary between more mobile (e.g., generalists) and sedentary ecomorphs (e.g., head). Wing lice, in some cases, use phoresis as a way of moving between the hosts; thus, it is likely that their leg morphology may affect their ability to grasp onto the flies. We performed two geometric morphometric analyses using the shape of the head, one with a newly constructed phylogenomic tree and one without, to determine if each ecomorph resides in its own geometric space to understand how similar selective pressures, resulting from escape from host defenses, have resulted in repeated morphological changes across these ecomorphs. Finally, we explore the morphospace of each ecomorph with respect to competition. We compare rates of evolution for lice that cohabitate on hosts with other ecomorphs against those who don’t. These results will demonstrate if signals of competition, divergence, and convergence can be identified.

## RESULTS

### Chewing Muscles

We found differences in the chewing muscles between ecomorphs, with head lice having the largest and wing lice the smallest proportional mandibular muscularization (Fig. 1). Chewing muscle proportional volume was significantly associated with ecomorph (KW test, H = 25.324, df = 3, p<0.001), whereas the sex of the specimen did not have a significant impact (H = 0.283, df = 1, p>0.05; fig. S1). Significant group-wise differences were found between head and wing ecomorphs, and between head and generalist ecomorphs (Dunn’s post hoc, p<0.001; with 95% confidence; Fig. 1 and fig. S2). When accounting for the phylogenetic non-independence of the data, we found that ecomorph still accounts for a significant amount of variation in the volume of the mandibular muscularization (p<0.01), with the same significant groupings head and wing, and head and generalist lice (p<0.01).

**Figure 1.**
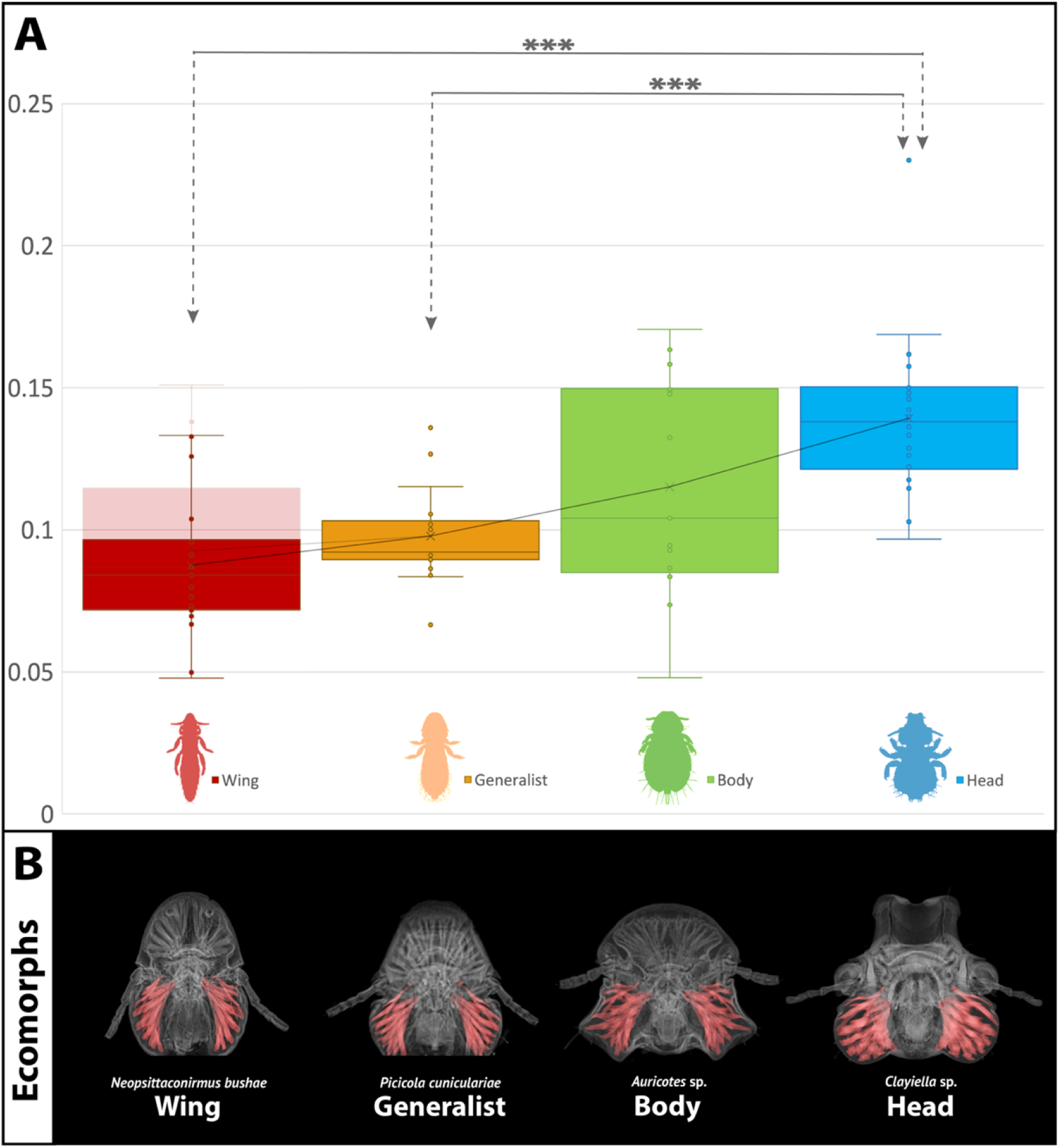
Mandibular muscularization of feather louse ecomorphs. (**A**) Proportional volume of the chewing muscles to the volume of the head (N=77 or 79 with Acidoproctus). Asterisks indicate significant differences between group comparisons (***p<0.001). Transparent color shows data including two Acidoproctus specimens, which have a strongly indented marginal carina between frontal projections, affecting the total volume of the head. (**B**) CT scan image of the head of four ecomorph groups with the mandibular muscles highlighted in red.

Two species from the wing louse genus *Acidoproctus* had the highest proportional volume of chewing muscles compared to other wing lice. *Acidoproctus* has an unusual head shape, with a strongly indented marginal carina between frontal projections, which reduces the total volume of the head and thus results in a higher proportional volume of chewing muscles. Thus, we also removed *Acidoproctus* species for comparison as it seems to be functioning differently from other wing lice (Fig. 1 and fig. S1).

### Legs

We found a significant difference in the proportional length of the legs between ecomorph groups, both when including all 6 leg segments or the whole leg combined (KW test, p<0.001) (Fig. 2). When including all 6 leg segments in a Dunn’s post hoc test, significant group-wise differences in proportional length were found between wing and generalist lice (p<0.01), and between both wing and body, and wing and head lice (p<0.001). The femur of the first pair of legs was not significantly different between ecomorphs (p>0.05). However, when focusing on whole legs (femur + tibia), all three legs were significantly different between ecomorphs (p<0.05) (figs. S3 and S4).

**Figure 2.**
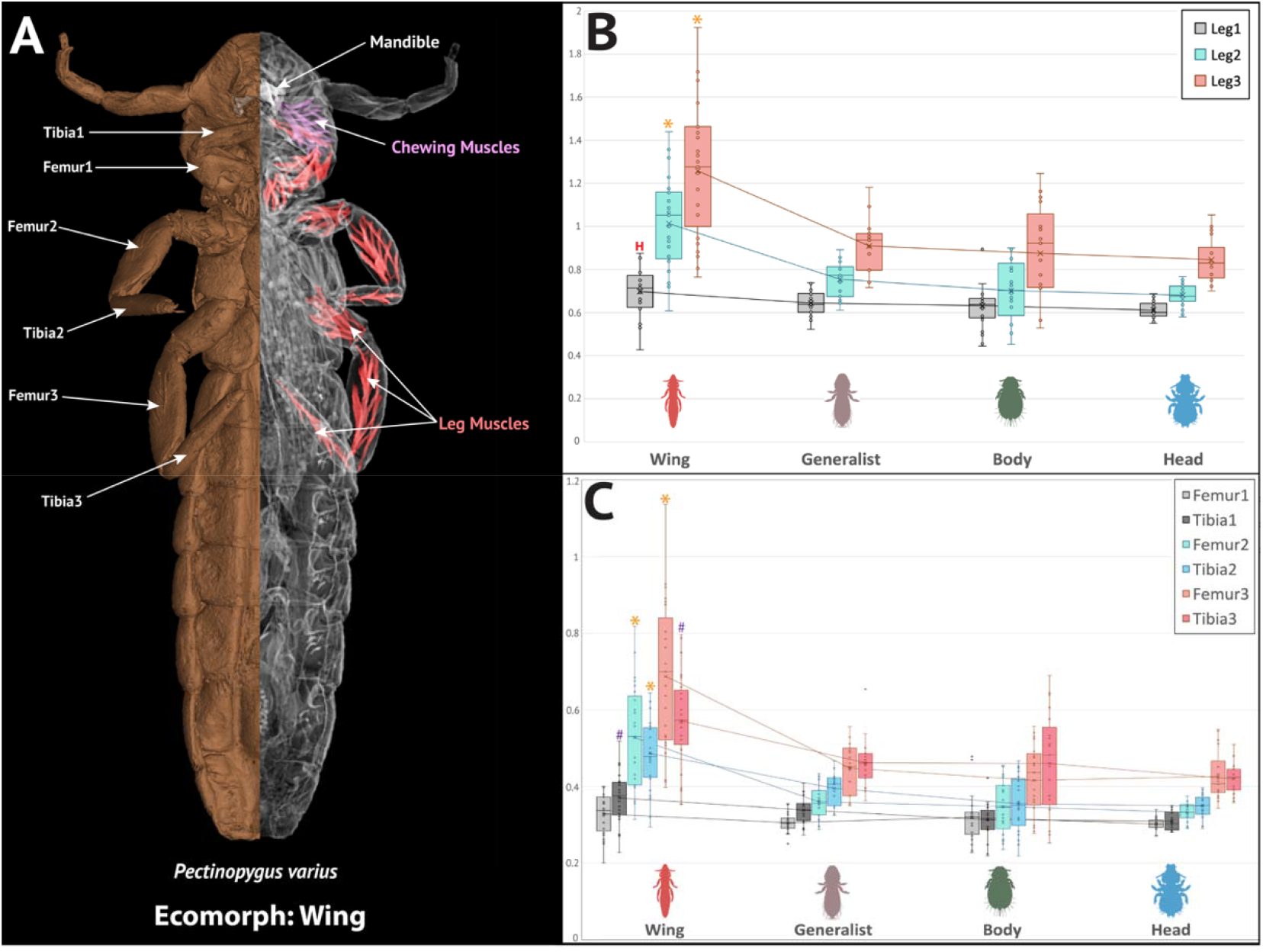
Proportional length of legs for each ecomorph group. (**A**) Illustrated louse morphology; (**B**) proportional length of legs (femur and tibia together) to metanotum width; (**C**) proportional length of leg parts (femur and tibia separately) to metanotum width. Asterisks shows the significant differences between wing ecomorph and all other groups; Hashes show significant differences between wing and both head and body groups; H letter shows significant difference between wing and head ecomorphs.

### Head Shape

The results of the principal components analysis (PCA) of the landmark data separated three of the ecomorph groups (wing, body, and head) into distinct regions of morphospace. Notably, generalist lice appeared in an intermediate region of morphospace between and overlapping each of the other three groups (Fig. 3). The greatest overlap was recorded between seven generalist specimens with five wing lice (Fig. 3). Nevertheless, some lice had a head shape more typical of a different ecomorph. For example, the wing louse *Aquanirmus occidentalis* had a head shape more representative of a generalist ecomorph. Conversely, the shape of the head of the generalist louse *Dahlemhornia asymmetrica* fell within the wing ecomorph morphospace. A similar scenario was found between head and generalist lice, where the generalist species *Brueelia antiqua* fell within the head ecomorph morphospace.

**Figure 3.**
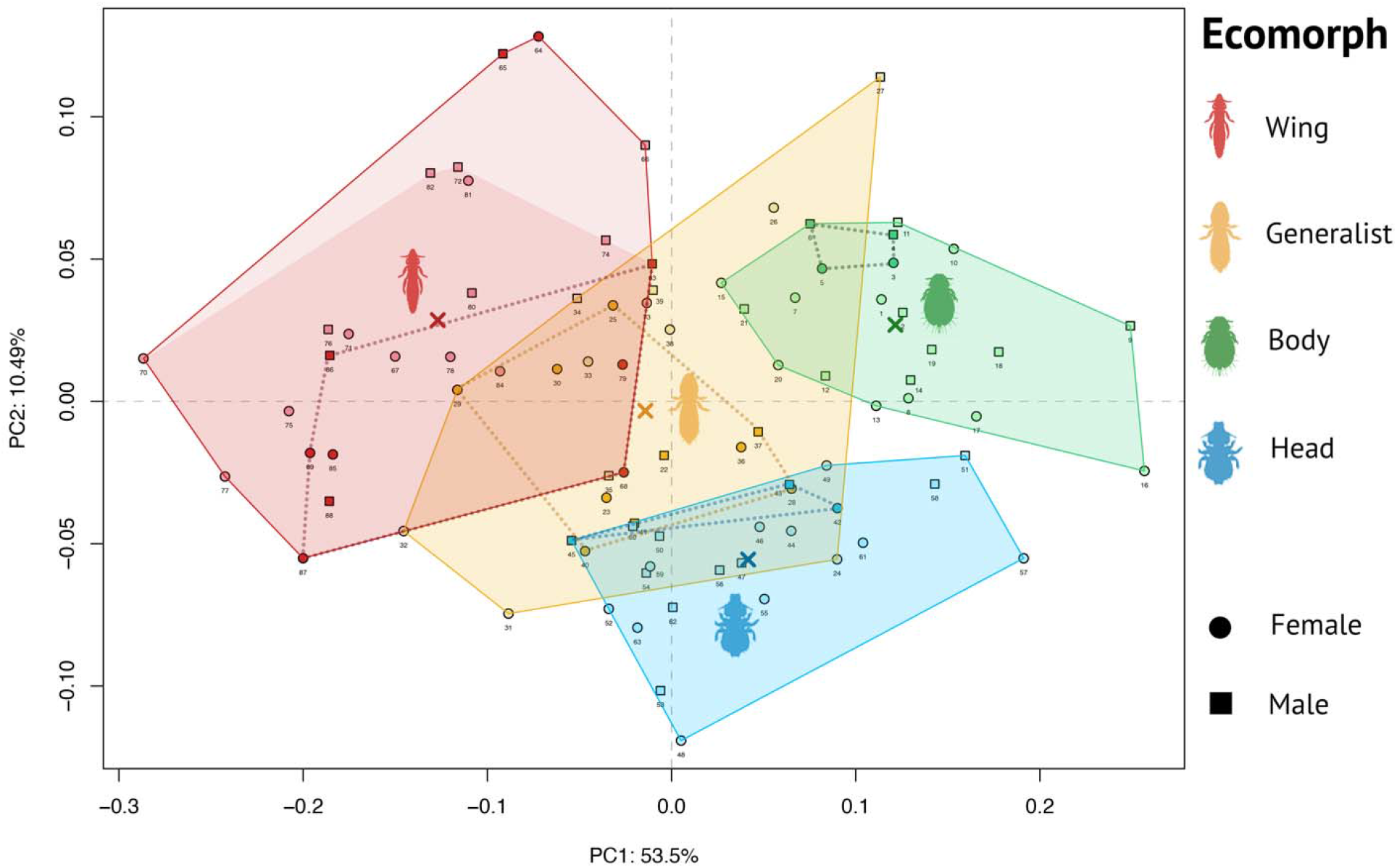
Feather louse morphospace. PCA results of generalized Procrustes analysis of 14 landmark data representing the head shape of 88 specimens belonging to four ecomorph groups (body, generalist, head, and wing). PC1(x) explains 53.5% of variability; PC2(y) explains 10.49%. The colored polygons represent the ecomorph group morphospace, with the mean PC1 and PC2 values for each ecomorph marked by an X. The brighter red background color polygon shows data including three Acidoproctus specimens (points 64–66), with the unusual head shape making their representation in the wing ecomorph group debatable. The dotted lines represent the space in which specimens (darker points) that do not cohabitate with other ecomorphs on their hosts are found. The points outside of those dotted lines indicate lice that can share their hosts with other genera of feather lice.

The mean PC value of the head ecomorphs was located in the overlapping space with generalist, while all other ecomorph groups had their mean values in their own discrete morphospace (marked with “X” in Fig. 3). This result suggests that head and generalist are the most similar ecomorphs. Interestingly, *B. antiqua* also had an unusually large mandibular muscularization (Fig. 3 and table S1). However, the remaining generalists that overlapped with the head ecomorph were only at the border of these two groups. There are few cases where the original qualitative assessments seemed to conflict with our major groupings. The specimen *Megaginus tataupensis*, which was originally assigned as a body louse (with other members of tinamou body lice), fell into the head louse morphospace. Similarly, *Bothriometopus macrocnemis*, originally characterized as wing louse, appeared within the body louse morphospace. Lastly, no generalists were found in the body louse morphospace. However, the head shape of two generalist specimens (female and male of *Colinicola mearnsi*) was unusual for a generalist ecomorph and more similar to body ecomorphs (points 26 and 27 seen in the top right corner of the PCA plot; Fig. 3).

Ecomorph type was also significantly associated with head shape when accounting for the phylogenetic non-independence of the data (phylogenetic ANOVA of PC1 and PC2, p<0.001). The significant group-wise differences found in PC1 were between wing and head, wing and body (p<0.001), wing and generalist, and body and generalist (p<0.01). Comparing PC2 (p<0.001), the significant group-wise differences were found only between head and all other ecomorph groups (p<0.01). Interestingly, while the head shape of ischnoceran chewing lice exhibited a significant phylogenetic signal, it was less than what was expected under a Brownian Motion model of evolution (K_mult_ = 0.69666; p < 0.001). Indeed, this result is congruent with the topology of the phylogenetic tree (Fig. 4), in which there were (Fig. 4) multiple cases of sister taxa belonging to different ecomorph groups. Similarly, the results of our phylo-morphospace analysis, which combined the PCA with phylogenetic tree data, showed that sister taxa were often associated with different morphospaces (Fig. 5). Here, the phylogenetically-aligned PCA (PACA), especially when using generalized-least squares (GLS)-centering and estimation, appears to affect the ecomorph grouping the most (Fig. 5D). Lastly, while there were no significant differences in net rates of evolution of head shape between ecomorph groups (p>0.05), feather lice that shared hosts with other ischnoceran feather lice had significantly higher evolutionary rates than species that were the only feather louse on their hosts (evolutionary rate of cohabiting species = 3.413e-06, rate for solo species = 2.544e-06, p=0.0053). There was no significant difference in the rates of morphological evolution between cohabiting vs. solo lice within each ecomorph (p>0.05).

**Figure 4.**
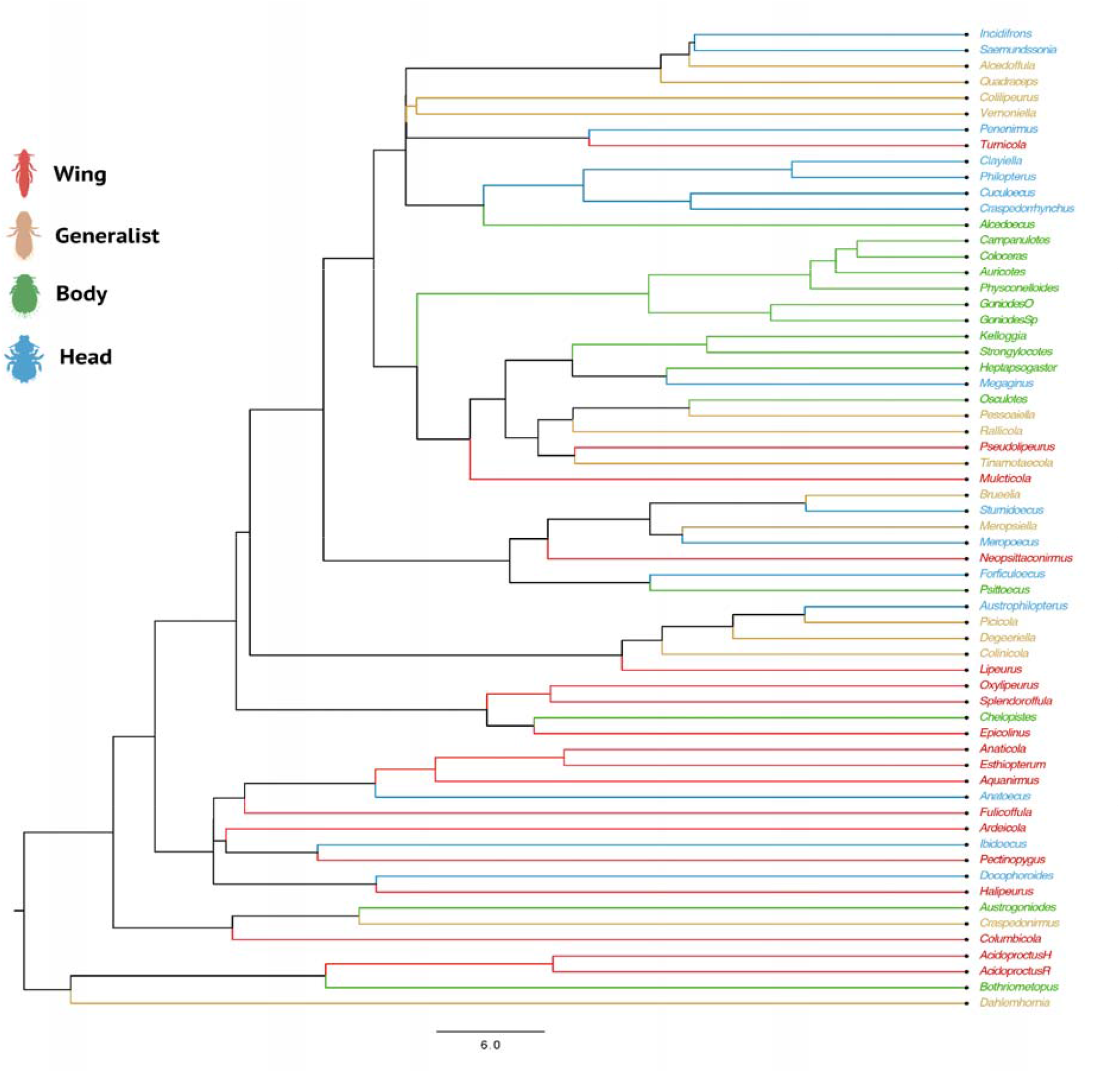
Phylogenetic relationships between feather louse genera belonging to four ecomorph groups. Specimens are colored according to their ecomorph identification.

**Figure 5.**
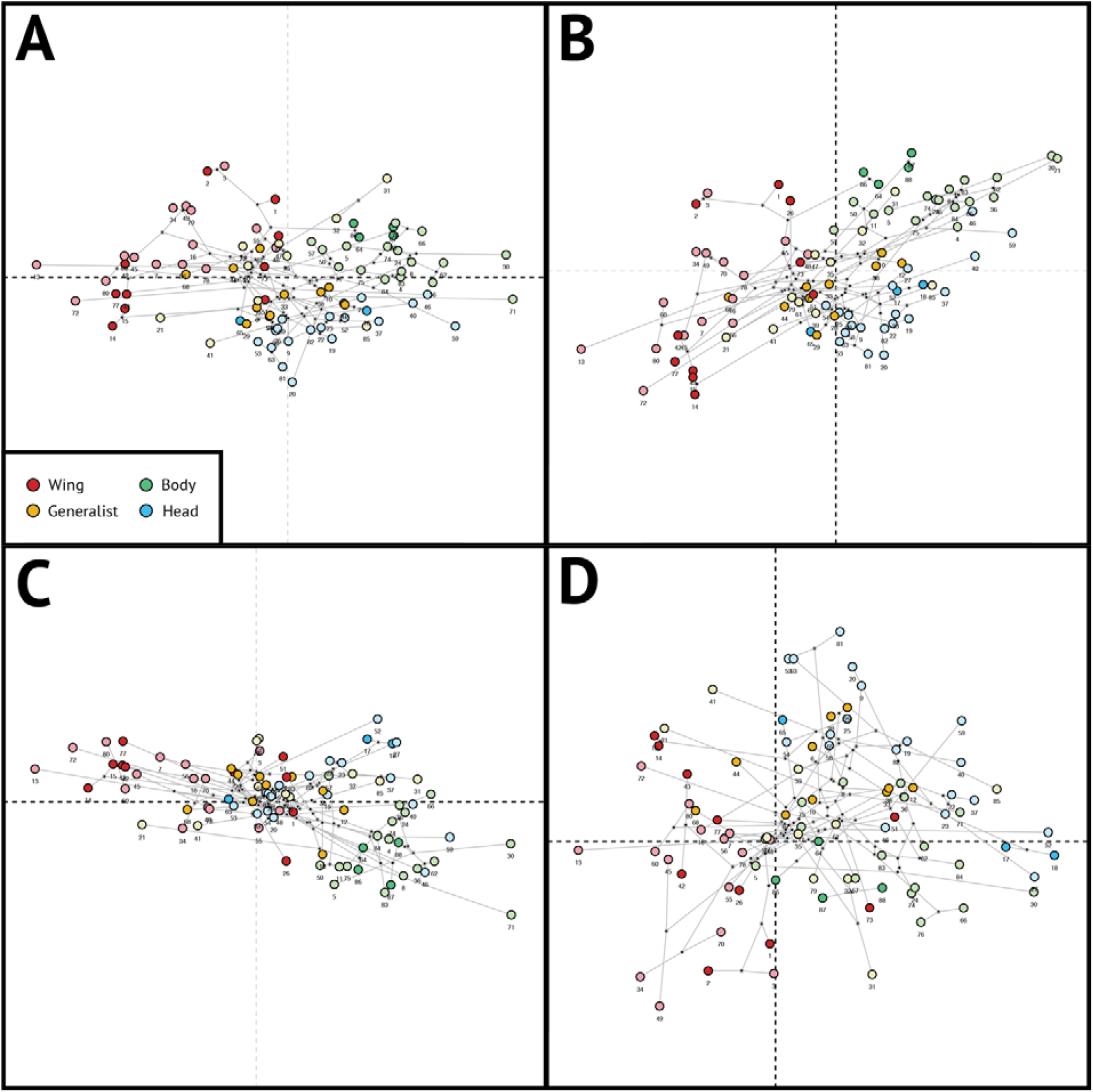
Phylogenetic (phy-PCA) and phylogenetically-aligned (PACA) principal components analysis of feather louse head shape. Darker colored points are specimens on host cohabiting with other louse species; light colored are species characterized as singleton feather lice. (**A**) phy-PCA using ordinary least-squares (OLS) centering and projection; (**B**) PACA using OLS; (**C**) phy-PCA using generalized least-squares (GLS) centering and projection; (**D**) PACA using GLS approach.

## DISCUSSION

Repeated convergent evolution in systems with similar selective pressures can help to illuminate the link between broad macroevolutionary patterns, ecological processes, and adaptations. Here we examined specific morphological changes between convergently evolved ecomorphs of avian feather lice and found that datasets from different morphological features have some degree of significant variation between the ecomorph groups. However, these features differ in which ecomorphs are significantly different, suggesting that different aspects of morphology are relevant to the specific adaptations of each ecomorph.

### Chewing Muscles

Head lice are thought to bite down to grip onto feather barbs to avoid being removed by scratching and, therefore, require larger chewing muscles than other ecomorphs. As predicted, we found that head lice have significantly larger proportional mandibular muscularization, followed by body, generalist, and wing lice (Fig. 1). The diet of all feather lice is very similar (*9*) in that all feather lice eat the downy portions of feathers, so it is reasonable to suggest that the major differences in chewing muscles are most likely related to escaping host defenses. In contrast to head lice, wing lice hide between the barbs of the wing feathers to avoid being removed by preening and do not grip these barbs with their mandibles, given the large size of the feather barbs in relation to louse size. To insert into this interbarb space, wing lice must be long and slender, and thus the size of their mandibular muscularization is likely constrained, because they would need expanded temples to accommodate larger muscles, as seen in head louse ecomorphs (e.g., Fig.1).

In mandibular muscularization, the most significant differences were found between head and wing and between head and generalist ecomorphs, and no significant differences were found between body lice and other groups. Body lice had the largest variance in the proportional volume of the chewing muscles (Fig. 1) and seem to be more variable in morphology than initially expected. This group likely needs additional attention to understand their full range of morphological variation and whether this relates to variation in escape strategy or to some other ecological difference. Additionally, as in the case of the genus *Acidoproctus*, an unusual head shape might be the cause of a higher proportional volume of chewing muscles. While this shows some limitations of our method, these cases are relatively rare, and, in general, using head volume to get a proportional volume of the chewing muscles appears to account for much of the variation in ecomorphology.

### Femur and Tibia Length

Limb length is expected to differ between ecomorphs due to their requirements for locomotion. Head lice are relatively sedentary, remaining mostly on the head, whereas generalist and body lice move around more freely on the host. Wing lice are the only ecomorph that move from the wing feathers to the body of the bird to feed on the downy feathers (*6, 20*). Focusing on the proportional leg length (femur and tibia separately or combined), the most variable group was the wing ecomorph (Fig. 2). Wing lice differed significantly from all other ecomorphs when we examined either femur or tibia separately (fig. S3) or combined (fig. S4). In contrast, body, head, and generalist lice feed on the down of the feathers in which they are located and escape from host preening in the same regions where they feed. This need for movement between the wing and body of the bird may explain why wing lice have legs with different proportions than other ecomorphs. In addition, wing lice are known to engage in phoresis on hippoboscid flies to disperse between hosts (*21, 22*). Harbison and Clayton (*23*) described how wing lice (*Columbicola columbae*) use their third pair of legs to grasp the fly’s leg and posited that their long limbs allow for a wider stance making phoretic dispersal possible (*23*). The high variance of leg length in our study confirms the likely importance of a larger proportional length of the second and third pair of legs in some species of wing lice (Fig. 2). For example, *Columbicola columbae* has a second and third pair of legs with a proportional value well above the average of the group (Fig. 2 and table S1). Body lice have not been found to use hippoboscid flies for phoresis (*23*). Some genera of head and generalist lice use phoresis (e.g., *Sturnidoecus* and *Brueelia*), but typically they attach to the fly with their mandibles rather than their legs (*6*).

To understand variation in leg morphology, focusing on single leg segments reveals the underlying differences. First, in the results of the proportional length of the whole legs (tibia + femur combined), the only difference from previous results of the whole legs was in the first pair, where the significant differences are found only between the head and wing lice. Second, when focusing on the proportional length of the separate leg segments, the femur of the first pair of legs does not significantly differ between ecomorphs. Additionally, the tibia length of the first and third pair of legs differs significantly between wing and body, and between wing and head ecomorphs, but not between wing and generalist. Both the tibia and femur of the second leg and femur of the third leg correspond to the differences between wing lice and all other groups.

While Harbison *et al*. (*21*) stated that both wing and body lice could initiate phoresis on flies, they found that wing lice had significantly lower detachment rates compared to body lice when flies were walking, grooming, or flying. In addition, body lice were found not to be attracted to hippoboscid flies, suggesting this is not a primary mode of transportation (*21*). Thus, the dominant variability in femur and tibia length between wing lice and all other ecomorph groups suggests their importance in prolonged attachment during the flight of hippoboscid flies. It is well documented that the long and slender shape of wing lice is an important strategy to avoid host removal by preening, in which lice insert between the barbs of the wing feathers (*15, 20, 21*). Our results suggest the potential importance of the length of the second and third pair of legs in dispersal.

### Head Shape

Geometric morphometric analysis of head shape reveals that head, body, and wing ecomorphs occupy distinct regions of morphospace, with generalist lice intermediate between these groups (Fig. 3). These results agree to some degree with a hypothesis that differences in the relative l volume of chewing muscles will result in different ecomorph head shapes, with the posterior part of the head (where the mandibular muscularization is located) becoming expanded in species with large chewing muscles. Interestingly, in the analysis of the proportional volume of chewing muscles, there was no significant difference between body and head louse ecomorphs. However, when accounting for the phylogenetic non-independence of the landmark data, head lice differ significantly in PC2 values from all other ecomorphs, including body lice (p<0.05). Thus, even while there was a high variance in chewing muscle volume in body louse ecomorphs, they still seem to occupy a morphospace distinct from head lice. Moreover, these results suggest that there are additional morphological differences in head features between the ecomorph groups other than chewing muscles. Lastly, most generalist lice occupy an intermediate area of morphospace to the more specialized ecomorphs, perhaps mirroring their ability to traverse the body and wing coverts. Interestingly, a few generalist species have unusual head shapes for the group to which they have been assigned. *Colinicola mearnsi* occupies a unique area of head shape morphospace, closer to the body louse morphospace. *Brueelia antiqua* is relatively close to the mean shape of the head louse (marked as blue “X” in Fig. 3) and has a strong mandibular muscularization. Interestingly, species of *Brueelia* are known for their phoretic capabilities on hippoboscid flies; however, they bite down on the fly to hold on rather than using legs like the wing lice (*24*). This aspect could explain their need for stronger mandibular muscularization for successful host switching. While the emu louse *Dahlemhornia* appears to be well within the wing morphospace, this genus has an asymmetrical head, potentially affecting the head shape analysis. These results suggest there are additional selective pressures on these shapes that warrant further study.

As previously mentioned, body lice have a high variability of chewing muscle volumes, and, similarly, there appear to be at least two different groupings of head shape in this group. One of these are “typical” body lice with a well-rounded head, e.g., the genus *Goniodes*. Another group of “atypical” body lice includes the genera *Bothriometopus, Strongylocotes*, and *Kelloggia*, which have head shapes more reminiscent of those from the generalist group. However, these genera have a strong mandibular muscularization (coefficient value 0.13–0.17) well above most generalist lice (0.08–0.11) (table S1). While *Bothriometopus* was initially assigned to the wing louse category (*25*), it falls well within the body louse category in all our analyses, with stronger mandibular muscularization, wider temple region of the head, and with the femur of the third pair of the legs being shorter than the tibia. In contrast, wing lice tend to have lower mandibular muscularization and the femur of the third pair of legs is usually longer than the tibia (table S1). The only exception here is the wing louse genus *Acidoproctus* which, interestingly, is the closest relative of *Bothriometopus* (Fig. 5), both of which can be found on waterfowls, order Anseriformes (*26, 27*). *Strongylocotes* and *Kelloggia* are tinamou lice members of the Heptapsogasteridae (*28*). This morphologically diverse group was generally assumed to be composed of body lice but might contain more variability than previously expected. For example, another tinamou louse in Heptapsogasteridae, *Megaginus tataupensis*, was initially assigned to the body louse category (*25*). However, our analysis suggests that this species is more appropriately assigned to the head louse group. The PCA morphospace analysis placed it well within the head louse space (Fig. 3), and it has strong mandibular muscularization (Fig. 1). This genus also has a triangular-shaped head and a rounded body more typical of head lice. Thus, here we demonstrate the value of a quantitative method for identification of ecomorph assignment in the future. Future analyses should also incorporate body shape, which is also a potentially important morphological feature separating feather louse ecomorphs.

### Phylogenetic Structure of Ecomorphs

By taking into account the phylogenetic relationships between these taxa, we were able to test whether the shape of the head is more similar among closely related vs. more distantly related lice from the same ecomorph. The phylogenetic ANOVA suggested that the type of ecomorph has a more significant effect on the shape of the head than evolutionary history. Additionally, while head shape in ischnoceran feather lice also exhibited a significant phylogenetic signal (p<0.001), closely related species were less similar to one another than what was expected under a Brownian Motion model of evolution (K_mult_<1). This result can also be seen in the phylogenetic tree (Fig. 4), and the phylo-morphospace analysis, which combines the PCA with the phylogenetic data (Fig. 5). In all cases (Fig. 5, A to D), sister taxa can be found belonging to different morphospaces. Herein, when using PACA with GLS-centering and covariance estimation for phylogenetically-aligned components analysis of shape data (Fig. 5D), the various ecomorphs, rather than occupying a well-identified and distinct morphospace, appear to be more scattered across the ordinary plot. Considering that there is no significant difference in net rates of evolution between the ecomorph groups in head shape, this pattern indicates a process of repeated adaptive divergence of these parasites within host groups, but convergence when comparing parasites across host groups.

### Why are there different feather louse ecomorphs?

Experimental transfer experiments placing lice on novel hosts of different sizes have demonstrated that lice have a reduced ability to escape from host preening when on a host of a different size than their usual host, suggesting louse body size is under selective pressure (*15*). This selection can lead to rapid changes in louse body size, potentially leading to reproductive isolation over several generations (*29*). Other studies have also shown rapid evolution of the coloration of lice in response to host defense. Bush *et al*. (*14*) investigated crypsis in feather lice and found that lice living on areas of the bird’s body that the host can see (e.g., wing) have evolved background matching coloration to avoid being seen by the host. Other experimental transfer experiments have also demonstrated the rapid evolution of coloration by placing lice on light and dark hosts for a short period of time (*30*). In light of these experiments, it is reasonable to suspect that morphological adaptation to escape from host defenses could evolve relatively rapidly, depending on the part of the host’s body where this escape occurs. For example, wing lice transferred from large pigeons to smaller doves move onto the head to escape from host preening because they are too large to fit between the smaller interbarb spaces of smaller doves (*31*).

Interspecific competition might be another factor playing a role. For example, Cicchino and Valim (*32*) compared louse oviposition between host species infected with single and multiple louse species. All three louse species laid their eggs on the head or neck of the bird when there were no other species present. However, when more than one species of louse was present on the same bird, each species laid their eggs in a more limited space, and the eggs of two different species were never found on the same feather in this case (*32*). This behavioral flexibility of lice, either in the presence of host defenses or competing louse species, could drive lice into different microhabitat niches. Given that there are likely to be a limited number of microhabitats and mechanisms to escape from host preening on an individual bird, there is likely a relatively small number of ecomorphological types into which lice can diversify, leading to widespread convergence across clades.

This is in agreement with our results, where the head shape of lice that are known to share their hosts with other ecomorphs tend to be more morphologically divergent (e.g., found in more disparate parts of morphospace (Fig. 3), whereas those lice found as singletons (alone) on a host represent morphological space usually closer to the centroid point. More interestingly, there appears to be a pattern, where feather lice that cohabit with other feather louse genera have significantly higher rates of morphological evolution than species that are known to be the only feather louse on that host group. While the mean values of net rates of head shape evolution show the same pattern in both cases, when comparing only cohabiting vs. alone or when accounting for this difference between all ecomorph groups, there still appear to be some overlapping values, whose variance is as of yet unexplained.

### The advantages and disadvantages of nano-CT when compared to 2-D microscopy

Nano-computed tomography has proved to be a useful tool for quantifying morphological differences in very small insects, which are hard or impossible to obtain using traditional microscopy. While shape variation analysis can be traditionally performed using 2-D, slide-mounted specimens, the 3-D datasets allowed us to record differences in depth, as well as allowing us to view the specimen from any angle and conduct the linear measurements – such as those of leg segments – that would otherwise be challenging with a slide mounted specimen. While it is possible that the heads of unmounted specimens may be more easily deformed, when compared to a slide mount in which the specimen can be pressed flat, a complete study comparing these two datasets (2-D and 3-D) is necessary to fully understand the impact of preservation methods on head shape analysis of feather louse ecomorphs.

The biggest benefit of CT is the ability to recover continuous trait data – lengths, shape components, or volumetric data – of internal structures. While the relative resolution of the CT datasets has a clear impact on the accuracy of the results, by standardizing the volume of the chewing muscles against the volume of the head we were not only able to account for variation in the body size of the lice (which can differ substantially even between the lice of the same ecomorph group) and also minimize issues with the varying quality of data between scans. This means that lower quality data are expected to have both lower volumetric values of the specific structures (here the chewing muscles) and lower values of the head volume, resulting in a similar coefficient values. However, as the accuracy of CT scanning improves, we should be able to obtain higher-quality data more easily.

Summarizing all of our results, it appears that the convergence in overall morphology, focusing here on head shape, mandibular muscularization, and variation in the length of the tibia and femur, is the result of the same specific morphological transitions for different lineages of the same ecomorph. However, collecting more data, such as shapes and measurements of other body parts, the volume of mandibles, and other features, is necessary to explore this hypothesis further.

## MATERIAL AND METHODS

### Data collection & 3-D scanning

We sampled 89 specimens representing 62 species of lice from all four ecomorphological groups (table S1). Initial ecomorph assignment followed previously published categorizations (*7, 12, 25*). Specimens had been previously collected from avian hosts using either the postmortem ethyl acetate fumigation and ruffling method or dusting with pyrethrum powder for live birds (*33*) and stored in 95% ethanol at -80°C. We performed high-resolution X-ray computed tomography (CT) scanning at the University of Florida’s Nanoscale Research Facility using Phoenix v|tome|x M (Waygate Technologies) and Xradia Versa 620 (Carl Zeiss optics) nanotomography systems. In order to retain structural integrity of the mounting system while minimizing X-ray attenuation through excess or dense material, specimens were gently affixed to the surface of a length of low-density adhesive tape, which was then rolled into a tube and attached to a 3.5mm diameter carbon fiber rod. This technique allowed up to six specimens to be mounted at once and scanned in sequence using the batch function of the Phoenix system and multiple recipes for the Versa. X-ray tube voltage was kept low (40-50kV) to improve contrast, and geometric magnification (Phoenix) and a combination of geometric and optical magnification with a 4X objective (Xradia) was employed to minimize the voxel-size and facilitate the recovery of microanatomy. Following scanning, the samples were gently removed from the adhesive tape by submerging them in alcohol. Radiographs were converted to tomograms using filtered backprojection, with Datos|X R (Waygate Technologies) and Reconstructor Scout-and-Scan Control System (Zeiss) software.

CT scans were processed with VGStudioMax v3.5.5, using specific gray values of density to create 3-D voxel regions of (1) the full body, (2) the head, (3) the chewing muscles of the specimens to obtain their volumetric (cubic) values (n = 79 specimens), and (4) lengths of leg segments. 3-D shape files of the head were saved as mesh files in the community standard Polygon File Format (.ply) for geometric morphometric analysis (n = 88 specimens). The files were then downsized in MeshLab v20.7 using the Quadric Based Edge Collapse strategy. Finally, we collected surface measurements to obtain the width of the metanotum, and the lengths of the femur and tibia for each leg (n = 87 specimens), one low-quality image was removed due to uncertainty in measurement precision.

### Volumetric and Measurement Data

The proportional volumetric data of chewing muscles were used to explore the differences between the ecomorph groups. To obtain the proportional values for each specimen, we divided the volume of the chewing muscles by the total volume of the head. Similarly, we used length values of both the femur and tibia individually and together and divided each by the width of metanotum to obtain proportional length values. To determine if the values were normally distributed, we used a Shapiro-Wilk test of normality and found that neither chewing muscle volume nor leg measurements (conducted for each leg individually) were normally distributed (p<0.05). Therefore, we selected the non-parametric Kruskal-Wallis (KW) rank sum test and Dunn’s post hoc test for the following analyses. We looked for statistically significant differences between the ecomorph groups and sex using a KW test. To identify statistically significant differences, we used Dunn’s test and 2-sided p-value with a Bonferroni correction using the R package “GmAMisc” and function kwPlot (figs. S2 to S4). Finally, results of volumetric dataset were compared with a Phylogenetic ANOVA, using phylogenetic information (below) and adjusted p-value using a Bonferroni correction for multiple tests.

### Phylogenetic information

We followed Johnson *et al*. (*34*) for genomic sequencing of lice. Specifically, for each louse species and 25 outgroup taxa (Anoplura, Trichodectera, Rhynchopthirina, and Amblycera) (table S2), we did single specimen-based genomic DNA extractions, prepared Illumina libraries, and sequenced them on an Illumina NovaSeq sequencer to achieve at least 30–60x coverage of the nuclear genome. We then pre-processed data using fastp v0.20.1 [phred quality >= 30, (*35*)] and deposited the raw reads in NCBI SRA (table S2).

We used aTRAM v2.0 (*36, 37*) to assemble a target set of 2,395 single-copy ortholog protein-coding genes, using a panel of amino acid sequences from the human head louse *Pediculus humanus* as the reference. After assembly, we removed intron sequences using an Exonerate-based (*38*) stitching pipeline (atram_stitcher) that identifies exon sequences and stitches them together (*39*). The whole assembly and phylogenomic pipeline, including processing steps, parameters, and commands, followed previous studies (*34, 40*).

We translated nucleotide sequences into amino acids and aligned them using MAFFT v.7.471 (*41*). After that, we back-translated sequences to nucleotides and used trimAL v.1.4.rev22 (40% gap threshold) to trim individual gene alignments (*42*). We discarded any gene that was represented by fewer than four taxa. We then concatenated gene alignments into a supermatrix and ran an IQ-TREE 2 v.2.1.2 partitioned analysis that included model selection for each partition (*43, 44, 45, 46*). We estimated support using ultrafast bootstrapping in IQ-TREE (*47, 48*). We rooted the tree on Amblycera based on prior studies (*8, 40*).

Lastly, we produced an ultrametric tree using the least square dating (LSD2) method implemented in IQ-TREE (*49*). For this analysis, we used the same calibration points as in Johnson *et al*. [(*40*) root age 92 mya)] and set a minimum branch length constraint (u = 0.01) to avoid collapsing short but informative branches without introducing bias to the time estimates (see https://github.com/tothuhien/lsd2). For better visualization, we colorized the phylogenetic tree according to the ecomorph groups (Fig. 4). Additionally, we used the phytool function “bind.tip” in R (*50*), to split the tree tips to have both female and male represented and corresponding to the morphological dataset.

### Geometric Morphometric Analyses of 3-D Landmark Data

To analyze the head shape between the ecomorph groups, we used 3-D shape files in the R package “geomorph” (*51*), using our modified R script from Paluh *et al*. (*52*) (see https://doi.org/10.5281/zenodo.7557419 or https://github.com/StanleeKol/LouseEcomorphs). Fourteen homologous landmarks were digitized in shape files in the same order to best simulate the shape of the head (fig. S5, A and B). Then we turned these data into a 3-D array for a generalized Procrustes analysis (GPA; fig. S5C) to obtain shape variables from landmark data. Each point was rotated and aligned, and Procrustes coordinates produced using the geomorph package (https://CRAN.R-project.org/package=geomorph) (*51, 53*). Additionally, we visualized the estimated mean shape from our data (fig. S5D).

We ran a principal component analysis (PCA), and the Procrustes-aligned specimens were plotted in three dimensions of tangent space for 88 specimens (PC1, PC2; Fig. 3). We used our phylogenetic tree with split tips to account for both males and females (n=88) and ran the following analyses for 10,000 iterations. First, we ran a phylogenetic ANOVA to analyze the patterns of shape variation and covariation between ecomorphs in our set of aligned coordinates (running separately for PC1 and PC2). Next, we compared the net rates of morphological evolution of ecomorph groups and of cohabitating vs. alone species using simulation method for both. The function for permuting the Procrustes shape data among the tips of the phylogeny was used to identify phylogenetic signal using the K-statistic [K<1 less phylogenetic signal than expected; K>1 greater phylogenetic signal than expected under the Brownian Motion model of evolution (*54*)]. Additionally, a phylogenetic morphological PCA was conducted (Fig. 5), with estimated ancestral states and phylogenetic branches projected into ordination plots (*55*). We used two approaches: 1) a phylogenetic PCA (phy-PCA; Fig. 5, A and C), which reveals trends that are the most independent of phylogenetic variation; and 2) a phylogenetically-aligned PCA (PACA; Fig. 5, B and D), which reveals the variation that is most associated with the phylogenetic signal. Additionally, we used both variants for centering and projections via ordinary least squares (OLS; Fig. 5, A and B), and generalized-least squares (GLS; Fig. 5, C and D).

## Acknowledgements

We thank A. Hernandez and C. Wright at the University of Illinois Roy J. Carver Biotechnology Center for assistance with Illumina sequencing. We thank S. Virrueta Herrera for assistance with DNA extractions and K. K. O. Walden for assistance with submission of sequences reads to NCBI.

## Funding

This research was supported by the National Science Foundation under Grant No. DEB-1925312 to J.M.A., DEB-1924759 to E.L.S., and DEB-1239788, DEB-1925487, and DEB-1926919 to K.P.J., and the European Commission grant H2020-MSCA-IF-2019 (INTROSYM:886532) to J.D.

## Author contribution

Conceptualization and Investigation: S.K., E.L.S., K.P.J., and J.M.A. Methodology: S.K., E.L.S., A.P., and J.D. Visualization: S.K. Supervision: J.M.A. Writing— original draft: S.K., J.M.A., and A.G. Writing—review and editing: S.K., K.P.J., J.D., A.G., E.L.S., A.P., and J.M.A.

## Competing interests

The authors declare that they have no competing interests.

## Data and materials availability

All data needed to evaluate the conclusions in the paper are present in the paper or Supplementary Materials. Additionally, the R scripts and the data for head shape analyses are available at Zenodo (https://doi.org/10.5281/zenodo.7557420) and GitHub (https://github.com/StanleeKol/LouseEcomorphs).

## Notes

https://github.com/StanleeKol/LouseEcomorphs

### Competing Interest Statement

The authors have declared no competing interest.

