## Supplementary Figures S1-S5 for "Nano-computed tomography reveals repeated phenotypic divergence in parasites to escape host defense"

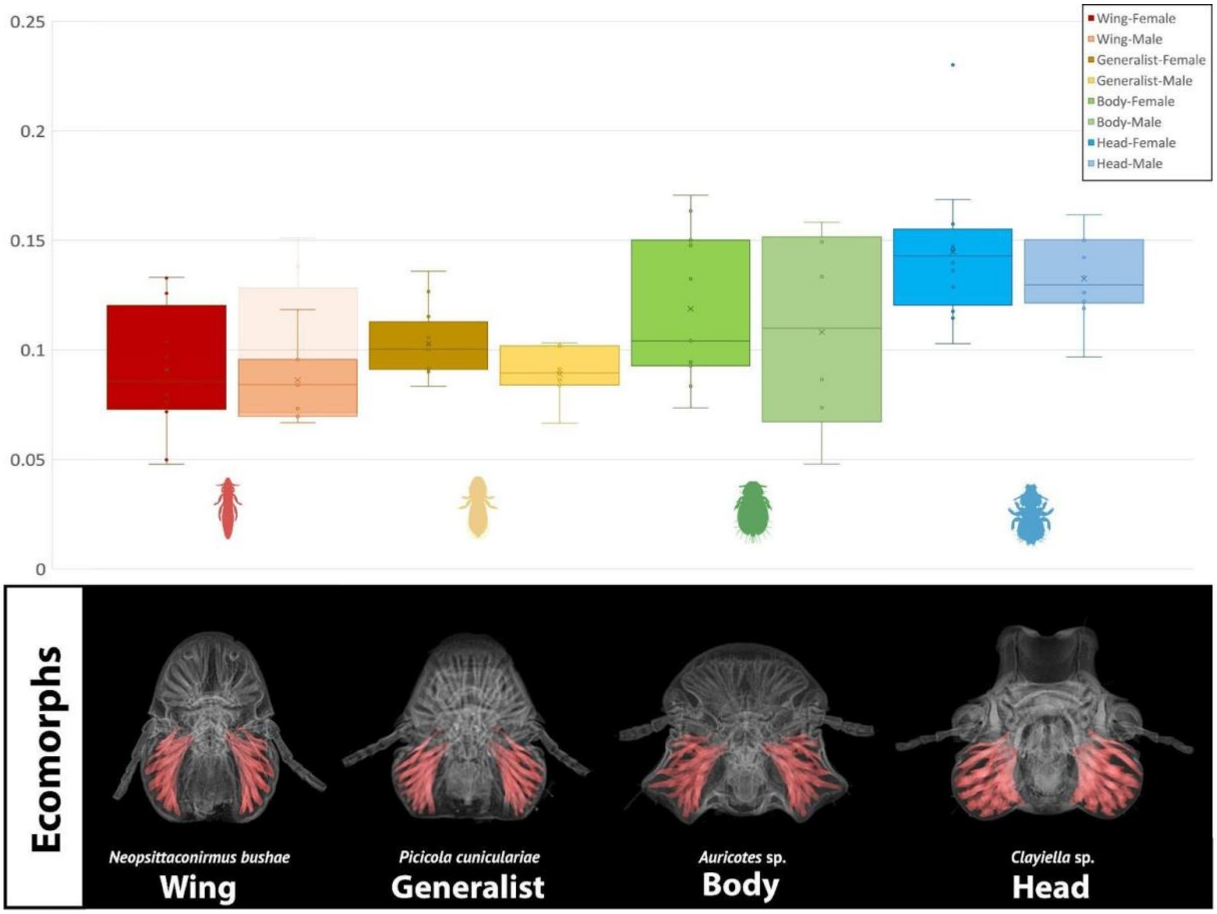


**Supplementary Figure S1.** Comparison of proportional volume of chewing muscles between feather louse ecomorphs and both males and females.

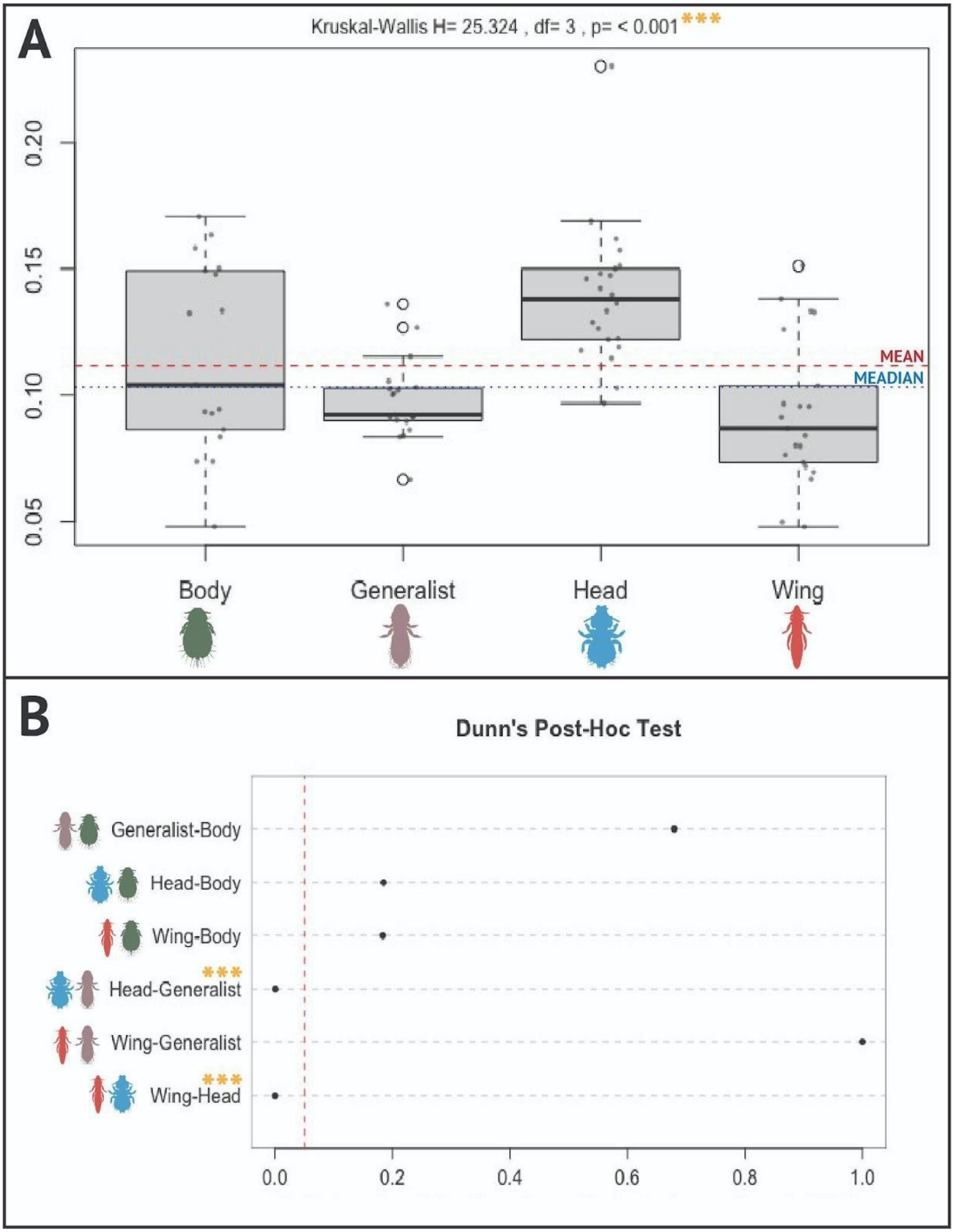


**Supplementary Figure S2.** Visualization of Kruskal-Wallis and Dunn's tests for chewing muscle proportional volume.

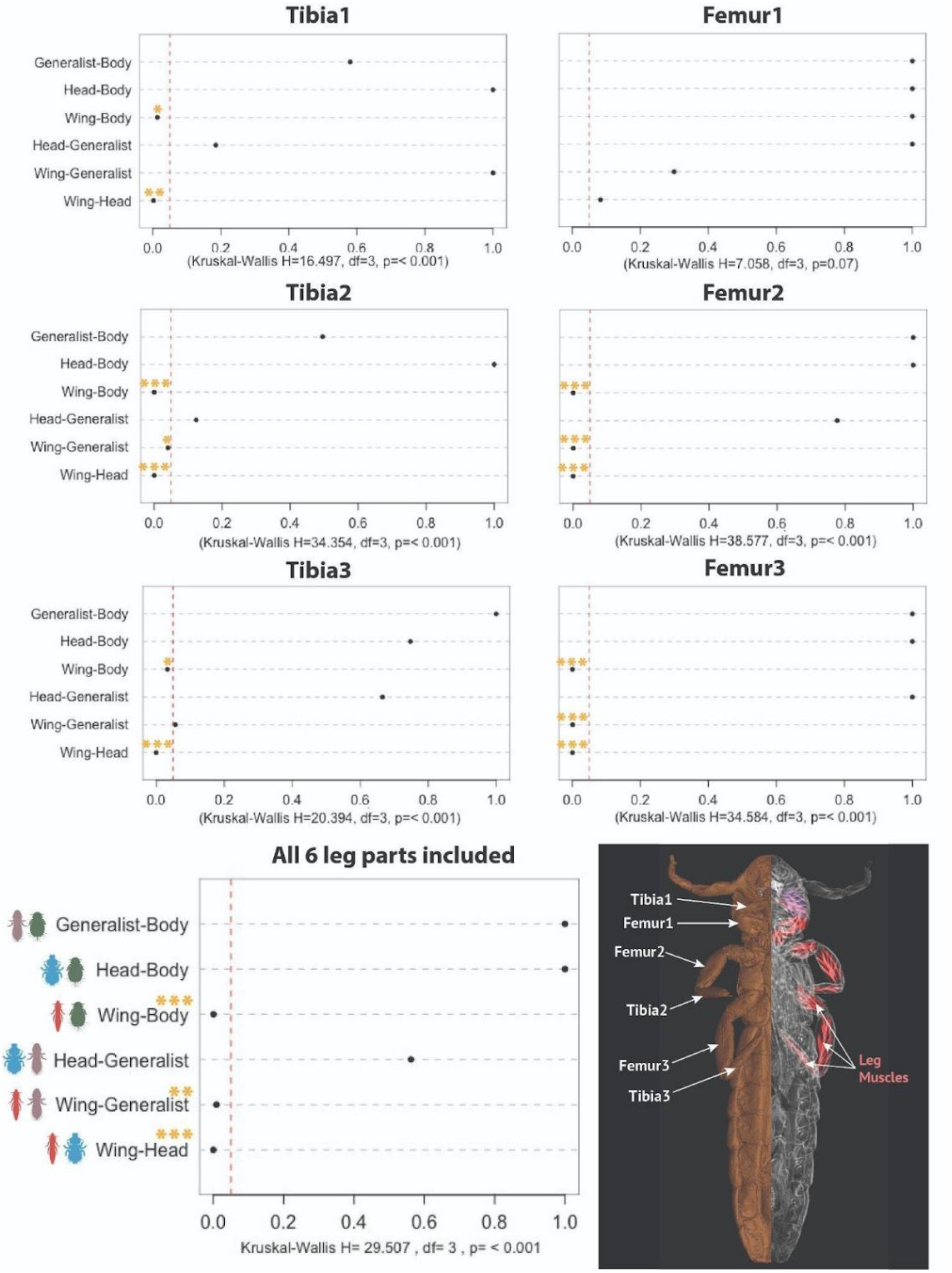


**Supplementary Figure S3.** Visualization of Kruskal-Wallis and Dunn's tests for proportional length of leg parts individually and all together (*p<0.05, **p<0.01, ***p<0.001).

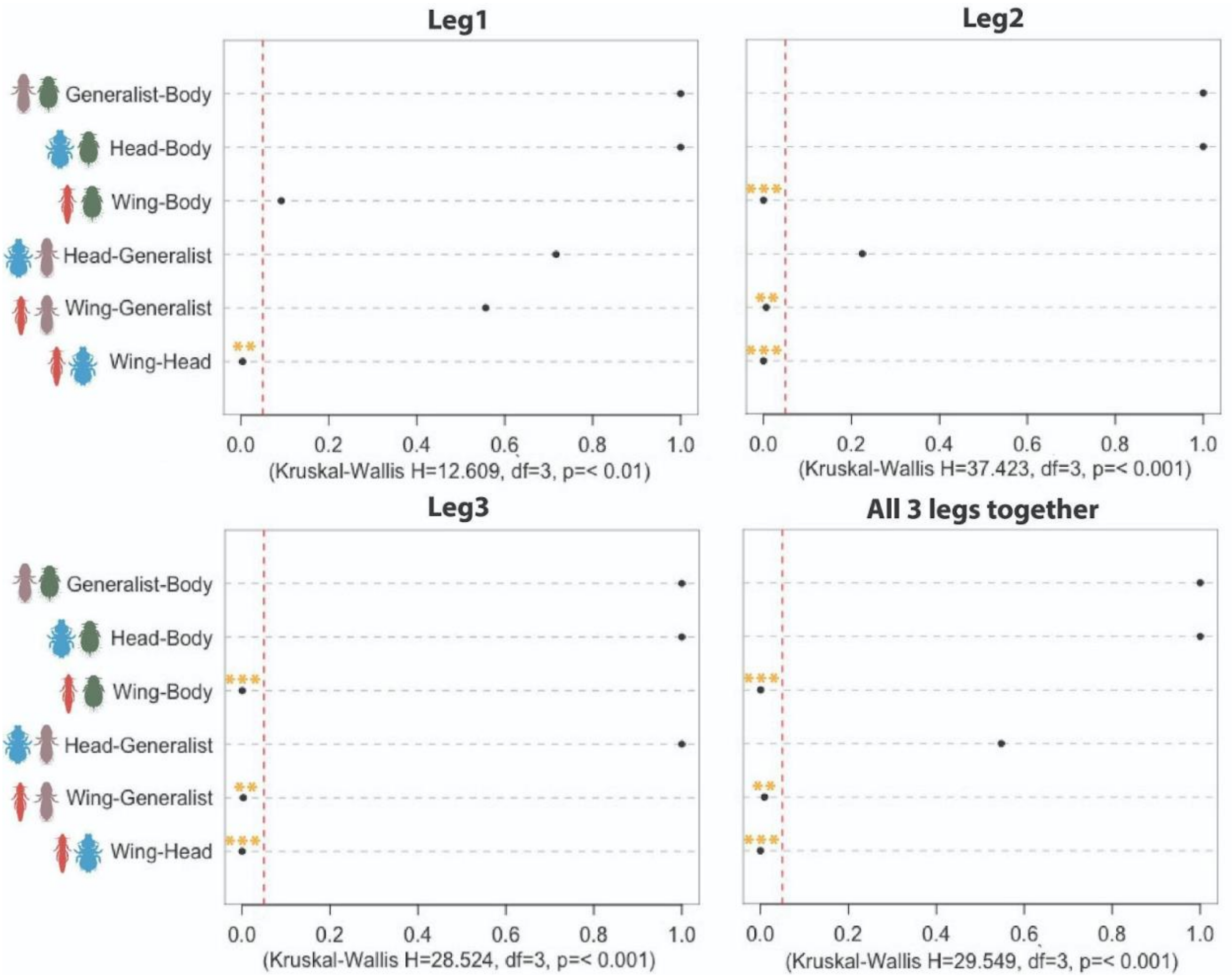


**Supplementary Figure S4.** Visualization of Kruskal-Wallis and Dunn's tests for proportional length of whole legs (tibia + femur) separately and as all three legs together (*p<0.05, **p<0.01, ***p<0.001).

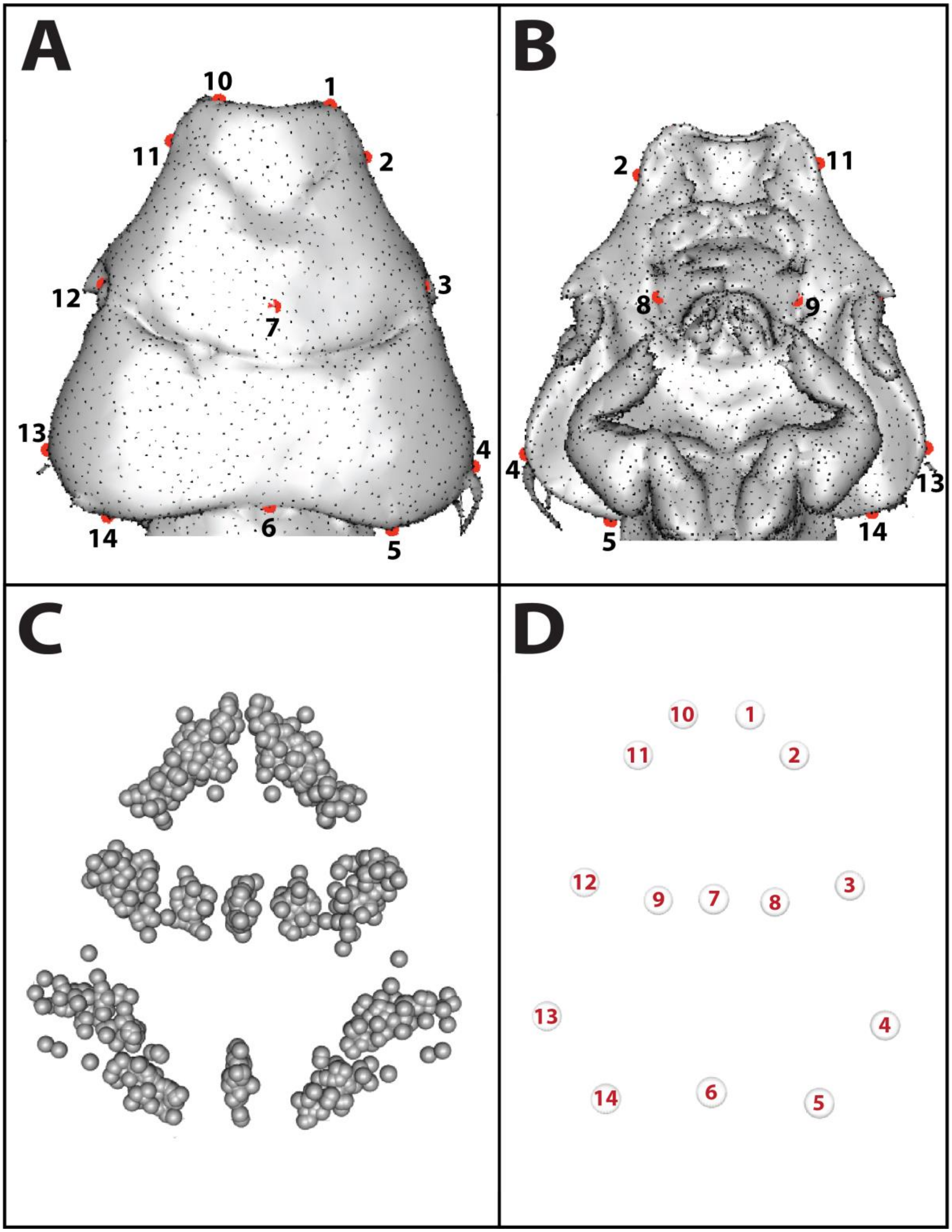


**Supplementary Figure S5.** Head shape data: a, b – 3-D landmarks used in our analyses; c – 3-D landmarks plotted after generalized Procrustes analysis (GPA); d – mean shape generated from the GPA data.

### Supplementary Tables

**Table S1:** List of feather louse specimens included in our analyses, with their ecomorph grouping, sex, voucher code, host species, country of collection, cohabitating status, and with Yes/No if used for landmark analysis (if yes, PCA labels are included), if used for analysis of proportional volume of chewing muscles (if yes, volume values of mandibular muscles, head, and their proportional coefficient are included), and if used for analysis of proportional leg length (if yes, metanotum width, leg length, femur and tibia length, and proportional coefficients values are included). N/A corresponds to missing data.

**Table S2:** List of louse specimens from which were obtained genomic data (including outgroups), with their ID codes, and Sequence Read Archive (SRA) accessions.
